# Comprehensive analysis of insertion sequences within rRNA genes of CPR bacteria and biochemical characterization of a homing endonuclease encoded by these sequences

**DOI:** 10.1101/2024.02.27.582315

**Authors:** Megumi Tsurumaki, Asako Sato, Motofumi Saito, Akio Kanai

**Affiliations:** Institute for Advanced Biosciences, Keio University, Tsuruoka 997-0017, Japan; Systems Biology Program, Graduate School of Media and Governance, Keio University, Fujisawa 252-0882, Japan; Faculty of Environment and Information Studies, Keio University, Fujisawa 252-0882, Japan

**Keywords:** Candidate Phyla Radiation, rRNA intron, homing endonuclease, bioinformatics

## Abstract

The Candidate Phyla Radiation (CPR) represents an extensive bacterial clade comprising primarily uncultured lineages and is distinguished from other bacteria by a significant prevalence of insertion sequences (ISs) within their rRNA genes. However, our understanding of the taxonomic distribution and characteristics of these ISs remains limited. In this study, we used a comprehensive approach to systematically determine the nature of the rRNA ISs in CPR bacteria. The analysis of hundreds of rRNA gene sequences across 65 CPR phyla revealed that ISs are present in 48% of 16S rRNA genes and 82% of 23S rRNA genes, indicating a broad distribution across the CPR clade, with exceptions in the 16S and 23S rRNA genes of Saccharibacteria and the 16S rRNA genes of Peregrinibacteria. Over half the ISs display a group-I-intron-like structure, whereas specific 16S rRNA gene ISs display features reminiscent of group II introns. The ISs frequently encode proteins with homing endonuclease (HE) domains, centered around the LAGLIDADG motif. The LAGLIDADG HE (LHE) proteins encoded by the rRNA ISs of CPR bacteria predominantly have a single-domain structure, deviating from the usual single- or double-domain configuration observed in typical prokaryotic LHEs. Experimental analysis of one LHE protein, I-ShaI from *Ca*. Shapirobacteria, confirmed that its endonuclease activity targets the DNA sequence of its insertion site, and chemical cross-linking experiments demonstrated its capacity to form homodimers. These results provide robust evidence supporting the hypothesis that the explosive proliferation of rRNA ISs in CPR bacteria was facilitated by mechanisms involving LHEs.

**IMPORTANCE:** Insertion sequences (ISs) in rRNA genes are relatively limited and infrequent in standard bacteria. With a comprehensive bioinformatic analysis, we show that in CPR bacteria, which are characterized by a high frequency of ISs, these ISs occur in 48% of 16S rRNA genes and 82% of 23S rRNA genes. We also report the systematic and biochemical characterization of the LAGLIDADG homing endonucleases (LHEs) encoded by these ISs in the first such analysis of the CPR bacteria. This study significantly extends our understanding of the phylogenetic positions of rRNA ISs within CPR bacteria and the biochemical features of their LHEs.

## INTRODUCTION

Culture-independent studies of microbial communities, such as metagenomic and single-cell genomic studies, have revealed numerous lineages that do not belong to known taxonomic groups (1). The Candidate Phyla Radiation (CPR) is a large monophyletic group composed primarily of uncultured bacteria (2, 3). The habitats of CPR bacteria are diverse, including fresh water (4), groundwater (5–9), subsurface sediments, hydrothermal vents (10), soil (11, 12), sludge (13, 14), and the human oral cavity (15), and only the phylum Saccharibacteria from the human oral cavity has cultured representatives (16). The CPR bacteria form a clade separate from all other bacteria on the phylogenetic trees constructed from 16S rRNA genes and concatenated ribosomal proteins (2, 3). CPR and other bacteria are also distinctly separated when clustered according to the presence or absence of specific protein families (17). In contrast, CPR is a sister group of Chloroflexi, part of the Terrabacteria group, but no conclusion has been reached regarding the phylogenetic status of CPR (18). Nonetheless, CPR accounts for a large proportion of the bacterial domain and cannot be omitted from the discussion of prokaryotic evolution.

CPR bacteria commonly have small cells and their genomes are often <1.5 Mb, and some strains are known to lack several genes encoding members of important metabolic pathways, such as the tricarboxylic acid cycle, amino acid biosynthesis, and nucleotide biosynthesis (6, 19, 20). Our recent studies have also shown that they lack several ribosomal proteins and some regions of rRNA genes (21). Such small cells and reduced genomes are commonly reported in known symbiotic or parasitic prokaryotes. Among the CPR bacteria, Saccharibacteria and some other strains are reportedly symbiotic with or parasitic on other organisms (16, 22–26), and although the biology of most other CPR bacteria is unclear, they are also thought to be dependent upon other organisms in some way.

Although CPR bacteria have reduced genomes, Brown *et al*. detected numerous insertion sequences (ISs) in their rRNA genes reconstructed from groundwater metagenomes (2). These ISs often have group-I- or group-II-intron-like sequences and protein-coding regions, which are removed from their transcripts. Group I or II introns have been found in eukaryotic, archaeal, and bacterial rRNA genes, although their occurrence in bacterial rRNA is limited to a few species (27). Group I introns have been found in the 23S rRNA genes of some thermophilic bacteria and obligate intracellular pathogens, and these introns are closely related to those in eukaryotic organelles (28–31). Introns in bacterial 16S rRNA genes are much rarer, and experimentally characterized group I and II introns have only been reported in giant sulfur bacteria (32). Conversely, some bacteria have intervening sequences (IVS) in their 23S rRNA genes, which are cleaved by RNase III without ligation, resulting in the fragmentation of the 23S rRNA. These are sporadically distributed among distant species and are thought to have been horizontally transferred (33–35). These IVSs are often found in parasitic species with reduced genomes, many of which have only a single copy of rRNA, indicating that ISs do not inhibit ribosome function. For example, rRNA ISs are conserved in all strains of *Coxiella burnetii*, which is an obligate intracellular pathogen (35). Although the biological role of rRNA ISs is unclear, it has been speculated that they confer some advantage in a symbiotic lifestyle (36).

Homing endonucleases (HEs) are often encoded in self-splicing elements, such as group I or II introns, and mediate “homing”, or the transfer of the intron to homologous intronless alleles (37, 38). HEs generate double-stranded breaks in their target sequences (approximately 10–40 bp), which are homologous to the insertion sites of host introns, leading to DNA repair by homologous recombination (39). HEs may also have a maturase activity that promotes IS splicing (40). There are several representative domain families of HEs, among which the LAGLIDADG and GIY-YIG families have been identified in CPR bacterial rRNA ISs (2). LAGLIDADG HEs (LHEs) are the best-studied HE family and are composed mainly of one or two LAGLIDADG domains. The single-domain LHEs function as homodimers (41, 42). Although the HE genes in introns are believed to undergo cycles of gain, degeneration, and loss (43, 44), it is unclear whether HEs encoded in CPR bacterial rRNA ISs are functional and retain their transferability or are remnants that have lost their function. The abundance of rRNA ISs in CPR bacteria is a noteworthy characteristic and is considered important in the evolution of rRNAs and introns. However, our knowledge of the phylogenetic distribution and functionality of the rRNA ISs in CPR bacteria remains limited.

In this study, we analyzed hundreds of rRNA gene sequences of CPR bacteria to detect and functionally classify their ISs. We found that rRNA genes containing group I or II introns are distributed widely across the CPR clade. We specifically report that single-domain LHE-like proteins are frequently encoded within these ISs. Importantly, we prepared the recombinant LHE-like protein I-ShaI, which is encoded by the rRNA IS of CPR Shapirobacteria derived from a high-temperature environment, and demonstrated biochemically that the enzyme has sequence-specific endonuclease activity and forms homodimers.

## RESULTS AND DISCUSSION

### Phylogenetic distribution of ISs in rRNA genes of CPR bacteria

To clarify the phylogenetic distribution of ISs in the rRNA genes of CPR bacteria in detail, we used a dataset of rRNA gene sequences (16S rRNA, n = 380; 23S rRNA, n = 348; 5S rRNA, n = 645) extracted from CPR bacterial genomes across 65 phyla (Table S1), which we obtained from public databases in our previous study (21). The 16S and 23S rRNA genes of CPR bacteria usually occur in only one copy per genome, with some exceptions. First, we detected the ISs of ≥100 bp by comparing the collected rRNA genes with those of *Escherichia coli* K-12. In total, 1346 ISs (378 in 16S rRNA genes and 968 in 23S rRNA genes) were identified, and the maximum number of ISs per gene was 16. No IS, as defined above, was found in any 5S rRNA gene. When the presence of ISs was evaluated for each gene, 183 of 380 (48%) 16S rRNA genes and 284 of 348 (82%) 23S rRNA genes contained ISs (Table S2). These proportions are significantly higher than the frequency of rRNA gene ISs in non-CPR bacterial genomes collected from RefSeq (1.1% for 16S rRNA genes and 18.5% for 23S rRNA genes). The maximum total length of ISs per gene was 5627 bp (mean 1075 bp, standard deviation [SD] 1043 bp) for 16S rRNA genes (n = 183) and 5750 bp (mean 1327 bp, SD 1112 bp) for 23S rRNA genes (n = 284). The ISs were distributed widely throughout the CPR bacteria but were not present in the 16S rRNA genes of Peregrinibacteria (Table S2). Although the IS estimated by comparison with *E. coli* rRNA do not correspond exactly to the regions removed by RNA splicing, it is obvious that the 16S and 23S rRNA genes of the CPR bacteria frequently contain ISs.

To characterize the rRNA ISs in CPR bacteria, we first detected the group I and group II introns (Table S3). Using the intron model from Rfam (45), 134 of the 378 ISs in 16S rRNA genes were classified as group I introns and 53 as group II introns; and 433 of the 968 ISs in 23S rRNA genes were classified as group I introns. No group II introns were found in the 23S rRNA gene. The open reading frames (ORFs) in the ISs were then extracted and annotated based on the Pfam protein domain database (46). The results showed that 455 ORFs (177 in 16S rRNA genes and 278 in 23S rRNA genes) encoded known protein domains, and that most of them encoded HE domains LAGLIDADG 1–3 or GIY-IYG (159 of 177 ORFs in 16S rRNAs, 274 of 278 ORFs in 23S rRNAs) (Table S4). Furthermore, the ORF encoding the 23S rRNA intervening sequence protein (23S_rRNA_IVP), which is a functionally uncharacterized protein domain known to encoded within the intervening sequences (IVSs) of known bacterial 23S rRNA genes, was detected in both the 16S and 23S rRNA gene ISs. That these protein families are encoded in ISs supports the results of a previous study (2). Here, the most frequently detected protein domain was LAGLIDADG_1, which accounted for 54% of the IS-encoded ORFs. ORFs encoding the LAGLIDADG_3 domain were only detected in the 16S rRNA gene ISs. The ISs were then classified based on the combination of group I or II introns and ORFs (Table S3). The number of ISs found to be group I or II introns and encoding annotated proteins was 124 in 16S rRNA genes (85 in group I and 39 in group II) and 211 in 23S rRNA genes (group I only). Moreover, there were 63 ORF-less introns in 16S rRNA genes (49 group I and 14 group II) and 222 in 23S rRNA genes (group I only). All the GIY-YIG family proteins were encoded by ISs that were not similar to either intron. The rest were uncharacterized ISs that did not match either intron and did not encode an ORF, most of which were relatively short (<200 bp), although long sequences (500–1000 bp) of unknown function were also detected. Some bacterial 23S rRNA genes are known to contain IVSs that are cleaved by RNase III without ligation, which are thought to be spread by horizontal transfer (34, 35, 47). Therefore, the CPR may also contain this type of IS.

To characterize the phylogenetic distribution of ISs in the CPR bacteria, the total IS length and the presence of group I or II introns and each protein family were summarized for each individual rRNA gene and mapped according to their phylum-level phylogenetic relationships (48). As shown in Fig. 1, group I or II introns and IS-encoded HEs are distributed widely throughout the CPR; however, as mentioned above, the 16S rRNA genes of Peregrinibacteria do not contain ISs, and in the closely related phylum Saccharibacteria, 64 of the 66 rRNA ISs were short sequences (≤250 bp) that did not contain group I or II introns or HE genes. Because many rRNA genes had multiple ISs, we investigated the positional distribution of the ISs in the genes according to their functional classification. First, we found that most of the ISs were concentrated at specific sites in the genes, and the frequent insertion sites were numbered 16IS1 to 16IS10 for the 16S rRNA genes and 23IS1 to 23IS14 for the 23S rRNA genes (Fig. S1). There was no relationship between the positions of ISs and the phylogeny of the host genome, and ISs were detected in a wide range of species at all positions. In the 16S rRNA genes, there were multiple insertion sites for group I introns (16IS4–7 and 10), whereas group II introns were concentrated predominantly at a single site (16IS9). In the 23S rRNA genes, the positions of ISs containing group I introns (23IS6–23IS14) were biased towards the 3′ region of the gene. Some of these group I or II intron insertion sites were consistent with known bacterial rRNA intron sites (28–32), whereas the group I introns at 16IS4, -5, -6, and -10 and 23SIS9, -10, -11, -13, and -14 have not been reported in any known bacteria. Many short ISs not classified as introns were found at sites 23IS2 and 23IS4, which are consistent with known sites of bacterial 23S rRNA IVS that are cleaved by RNase III, so they are probably the same types of IVS in the CPR.

**Fig 1.**
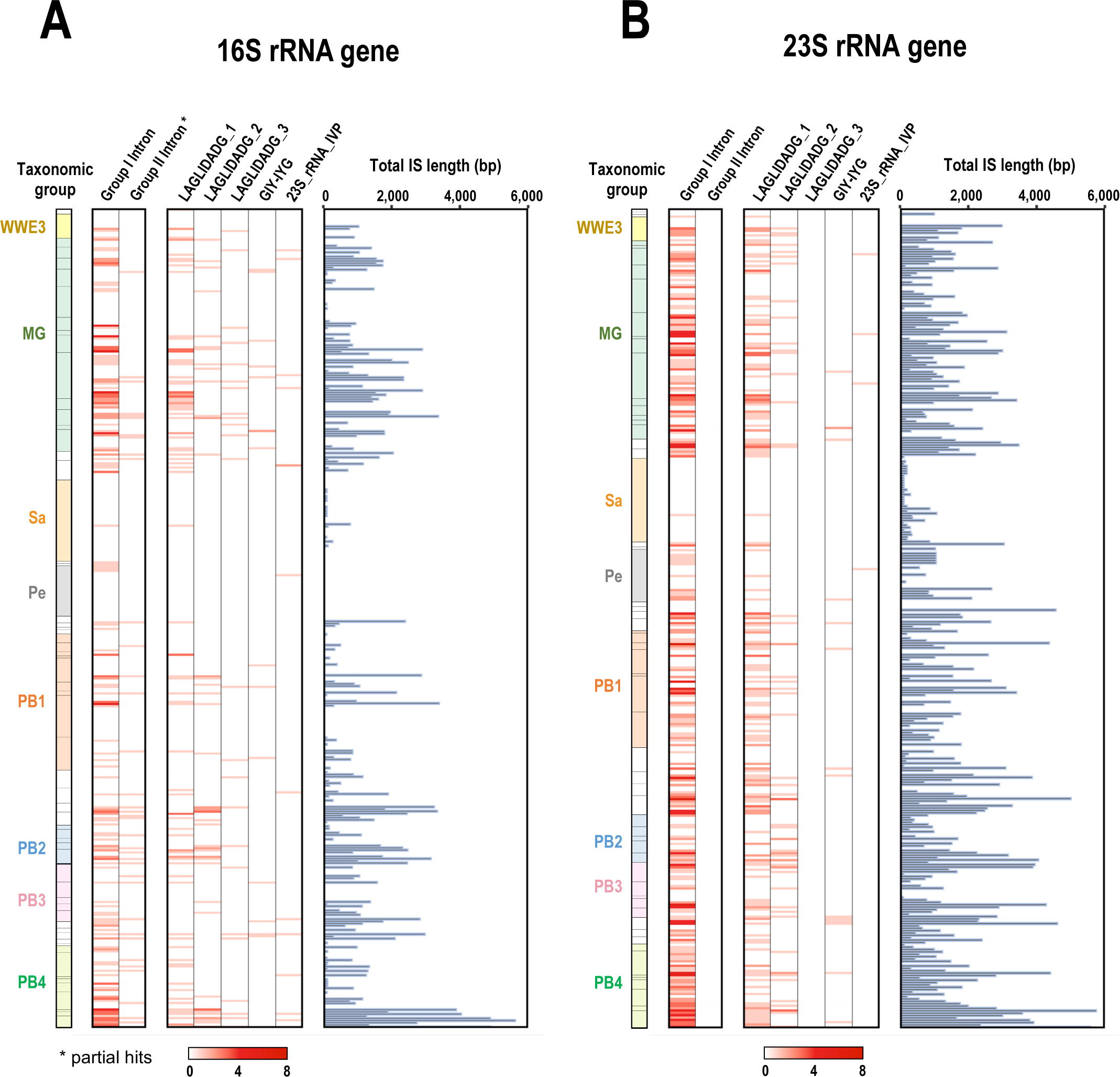
Distribution of ISs in rRNA genes according to the taxonomy of CPR bacteria. The presence and number of ISs that include introns and/or IS-encoded proteins and the total IS length are shown for each of the (A) 16S rRNA genes (n = 380) and (B) 23S rRNA genes (n =348) of CPR bacteria. The left panel shows the phylum-level taxonomy of CPR bacteria (48), and representative groups are colored (see Supplementary Table S1). MG; Microgenomates; Sa; Saccharibacteria; Pe; Peregrinibacteria; and PB; Parcubacteria.

In terms of ORF types, the ORFs located at the same insertion site roughly tended to encode the same protein family. In particular, the HE family LAGLIDADG was abundant at several specific sites. For example, the LAGLIDADG_1 protein was predominantly encoded in the ISs at position 1400 of the 16S rRNA gene (16IS9) and in ISs at positions 1917, 1931, 1952, 2593, 2604, and 2657 of the 23S rRNA gene (23IS6–8 and 12–14) (Fig. S2). The high density of LAGLIDAG_1 at these positions throughout the CPR clade means that it was probably acquired by the CPR ancestor and retained well throughout the divergence of the CPR, even if it was lost in some lineages. The LAGLIDADG_3 proteins, which were only found in 16S rRNA genes, were often located at the same sites as LAGLIDADG_2 proteins (16IS5 and 10). Both LAGLIDADG_2 and _3 proteins at these sites were randomly distributed in the host genome phylogeny, suggesting that they were not only derived by vertical transmission from the common ancestor but were acquired via multiple independent transfer events. Although these LAGLIDAD_2- and LAGLIDAD_3-domain-containing proteins did not have high similarity in the alignment, their structures predicted with AlphaFold2 were more similar to each other than either was to the LALGIDADG_1 structure (data not shown).

Although the LAGLIDADG proteins increased explosively at multiple sites, the GIY-YIG proteins, which are also members of the HE family, were encoded sparsely at several sites in the gene and did not appear to be increased at any specific site. This may be attributable to their relatively recent acquisition via horizontal transmission. 23S_rRNA_IVP were also scattered across several insertion sites, sometimes where other families predominated (16IS6, -7, and -9 and 23IS11). Among the ISs at the same insertion sites, such as 16IS4–7, 16IS 9–10, and 23IS6–14, some groups encoded LAGLIDADG proteins, whereas others were shorter and lacked ORFs. The distributions of both ORF-containing and ORF-less ISs were not clustered and were scattered throughout the host genome phylogeny. Furthermore, the alignment of these ISs revealed extremely high sequence similarity in regions other than the ORF. Some ISs contained degenerate ORFs with large deletions, suggesting scenarios in which only the ORF is lost from or inserted into the IS. ISs were found in both highly conserved and less conserved regions of the rRNA genes, but ISs encoding LAGLIDADG proteins (16IS4–7 and 9–10 and 23IS6–14) were only found in highly conserved regions, whereas other IS sites occurred in less-conserved rRNA gene regions. This is consistent with the assumption that LAGLIDADG was acquired from other organisms by target-specific homing. On the other hand, GIY-YIG seems to be outside the context of spread by homing, considering its low incidence in the CPR and low sequence conservation at its insertion sites. Most of the ISs were located at the top or base of the stem–loops in the rRNA secondary structures (Fig. S3). The ISs classified as group I introns (and often encoding LAGLIDADG) in the 23S rRNA genes clustered around the peptidyl transferase center in domain V, indicating that these ISs are correctly spliced and do not affect the activity of the ribosome.

### Homing endonucleases with a single LAGLIDADG domain discovered in rRNA ISs of CPR bacteria

LAGLIDADG is the most frequently encoded protein in the rRNA ISs of CPR bacteria. Of the 455 rRNA-IS-encoded ORFs containing known protein domains examined, 403 encoded LAGLIDADG family proteins (Table S4). To compare the characteristics of the LAGLIDADG proteins in CPR bacteria with those of other organisms, we collected a comprehensive set of LAGLIDADG-domain-containing proteins from the UniProtKB database. Sequence similarity searches using known LHE sequences as queries (Table S5) identified 5756 proteins, of which 5750 (eukaryotes: n = 4424; archaea: n = 109; CPR bacteria: n = 858; non-CPR bacteria: n = 356; viruses: n = 3) were obtained after those of unknown organisms were excluded. These proteins from UniProtKB, together with 377 LAGLIDADG family proteins extracted from rRNA ISs in CPR bacteria, were used for the following analysis.

The LAGLIDADG-domain-containing proteins in the CPR bacteria exhibited a different size distribution from those of the other taxonomic groups (Fig. 2). Specifically, whereas two size groups of approximately 180 and 310 amino acids were observed in non-CPR prokaryotes, the CPR bacteria contained few sequences with >300 amino acids, with peaks at around 180 and 210 amino acids. Eukaryotes showed a multimodal distribution, with a peak at around 310 amino acids, indicating the prevalence of proteins larger than those of prokaryotes. These differences in size distribution can be explained by variations in the protein domain structure. LHEs are typically composed of one or two copies of the LAGLIDADG domain (38). When we considered the domain structure of the LAGLIDADG proteins from UniProtKB (i.e., proteins from non-CPR bacteria) based on the Pfam database, sequences consisting of one or two copies of the LAGLIDADG_1 domain accounted for more than 70%, which were represented in the size distribution as peaks at around 180 and 310 amino acids (Fig. S4A). In contrast, the LAGLIDADG proteins from the CPR bacteria, with few exceptions, contained a single LAGLIDADG domain. Proteins with the LAGLIDADG_1 domain were predominantly around 180 amino acids in length, whereas those with the LAGLIDADG_2 domain were predominantly around 210 amino acids (Fig. S4B). Proteins with the LAGLIDADG_3 domain were detected in small numbers in archaea and CPR bacteria and were characterized by a non-domain region of variable length at the N-terminus. In addition to the typical one- or two-domain proteins, we also detected proteins composed of three or more LAGLIDADG domains, and proteins with LAGLIDADG domains in the intein regions (i.e., the regions excised during protein splicing) in other families of proteins (e.g., COX1, cytochrome B, etc.) (Fig. 3). These multidomain proteins were particularly abundant in eukaryotes and contributed to the formation of the multimodal size distribution.

**Fig 2.**
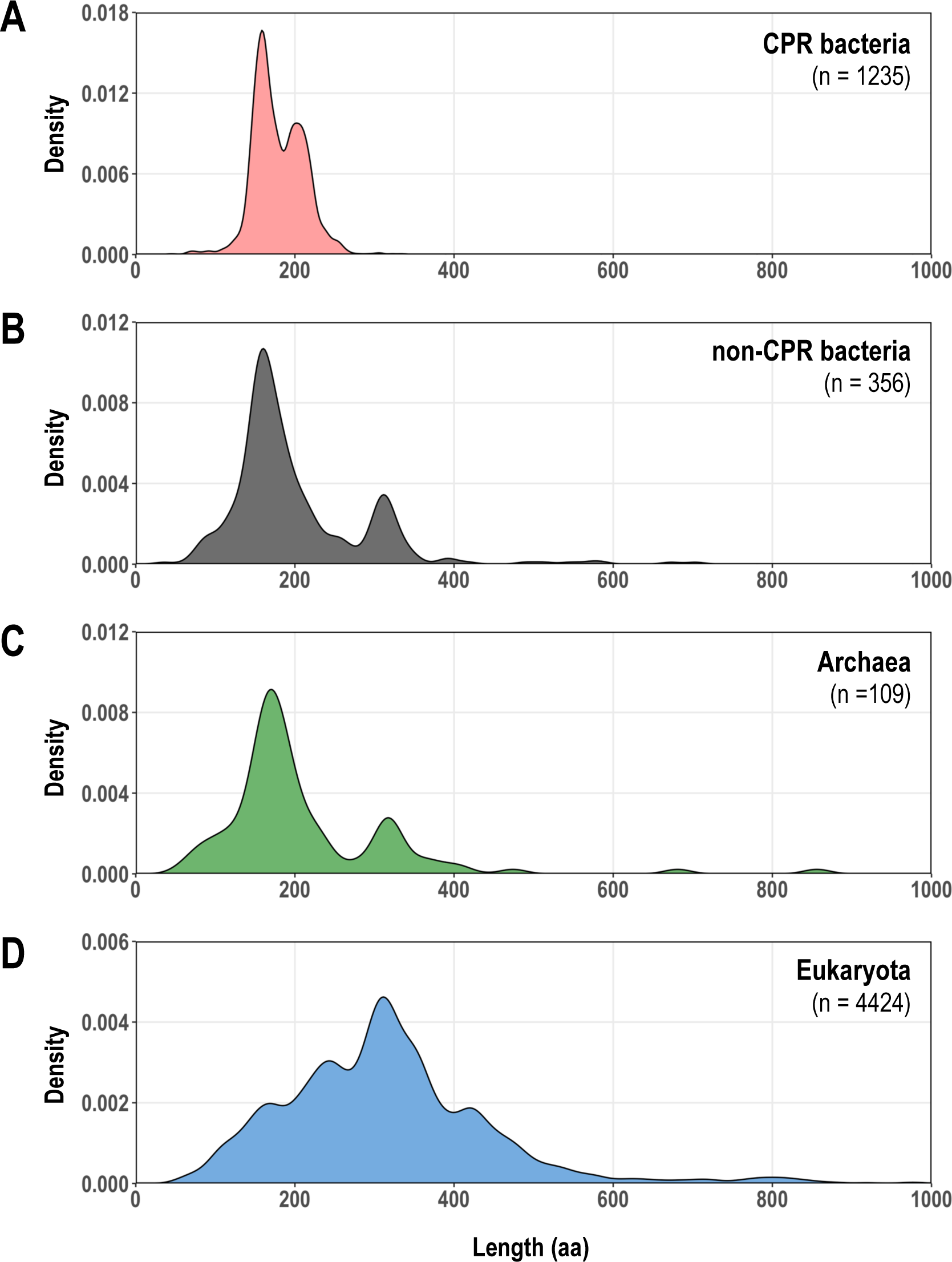
Comparison of the lengths of LAGLIDADG proteins encoded in CPR bacteria and other organisms. Length distributions of LAGLIDADG proteins are shown as density curves using the deduced amino acid (aa) sequences of the protein genes for (A) CPR bacteria (light red), (B) non-CPR bacteria (dark gray), (C) Archaea (green), and (D) Eukaryota (blue). Each horizontal axis displays only the length range over which the main peak is visible, excluding 18 proteins larger than 1000 aa.

**Fig 3.**
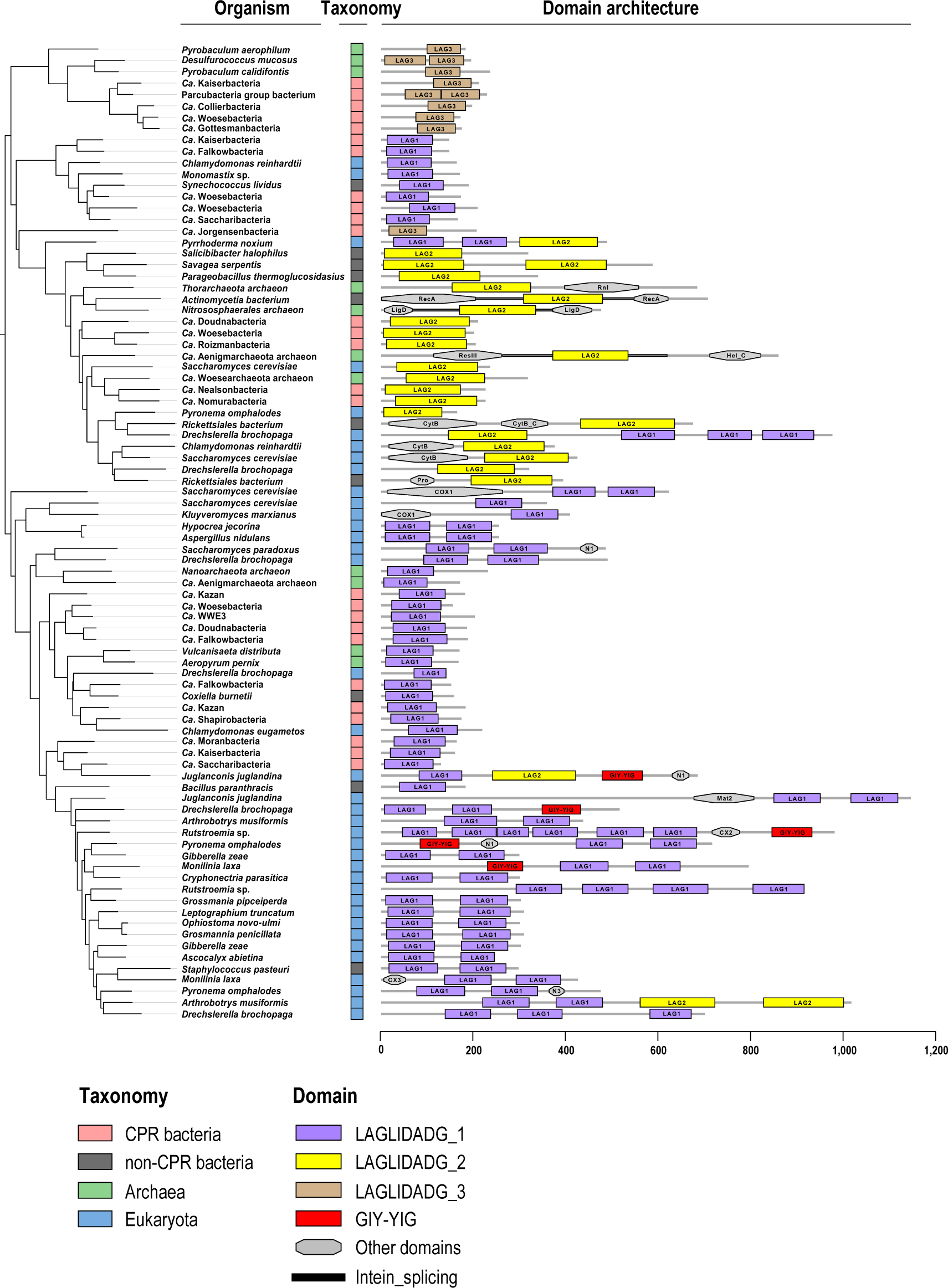
Examples of LAGLIDADG protein domain architectures. Phylogenetic tree of 86 representative LAGLIDADG proteins selected from Fig. 2 and their domain architectures are shown. The tree was constructed using the maximum likelihood method and rooted with midpoint rooting. Taxonomic groups of organisms are indicated by colored boxes: CPR bacteria, light red (n = 27); non-CPR bacteria, dark gray (n = 10); Archaea, green (n = 11); Eukaryota, blue (n = 38). The scale bar under the domain architectures shows the amino acid length of the protein. The domain name is based on the Pfam database: LAG1–3 (LAGLIDADG_1–3), Rnl (RNA_ligase), Hel_C (helicase_C), CytB (cytochrome_B), Pro (proton_antipo_N), N1 (NUMOD1), Mat2 (intron_maturase2), CX2 (COX2), and N3 (NUMOD3).

Phylogenetic trees based on the collected LAGLIDADG protein amino acid sequences showed that for each Pfam LAGLIDADG domain (LAGLIDADG_1, _2, and _3), the eukaryotic and prokaryotic sequences formed roughly separate groups (Fig. S5). Meanwhile, on the terminal branches, CPR sequences and non-CPR bacterial, archaeal, or eukaryotic sequences were sometimes very closely related, suggesting that the LAGLIDADG proteins have propagated among phylogenetically distant species. When a phylogenetic tree was constructed from only proteins containing a LAGLIDADG domain extracted from a CPR bacterial rRNA IS, the LAGLIDADG proteins encoded by ISs at the same gene position tended to be closely related to one another, even when the host genome lineages were distant (Fig. S6). When we considered the clades of LAGLIDADG proteins encoded at the same sites, some branches followed the phylogenetic relationships of the host genomes; however, in some cases, proteins derived from closely related host genomes occurred distantly within the clade, indicating horizontal transfer events caused by homing. The LAGLIDADG_2 proteins encoded in the IS at position 1498 of the 16S rRNA gene formed two distant clades, indicating that these were acquired in independent events. These results strongly support the proposition that the abundant rRNA ISs in CPR bacteria have not only increased by vertical inheritance from ancestral species but also by a site-specific homing mechanism mediated by the LAGLIDADG protein. Furthermore, there are examples of closely related LAGLIDADG protein sequences derived from different insertion sites, suggesting that ectopic transposition, as well as homing, has occurred.

### Biochemical characterization of LAGLIDADG protein I-ShaI derived from CPR bacteria

LAGLIDADG proteins encoded by rRNA ISs in CPR bacteria are thought to contribute to the propagation of rRNA ISs through a homing mechanism, but the activity of CPR-derived HEs has not been verified experimentally. Here, we examined the putative LHE encoded by *Candidatus* (*Ca.*) Shapirobacteria, identified as a CPR bacterium from the metagenomic-assembled genome of deep-sea hydrothermal vent sediments (with a sampling environment temperature of 48.5 °C) (10). We designated this enzyme “I-ShaI” in accordance with the established nomenclature (49). The enzyme encoded by the *I-ShaI* gene (DDBJ accession: BR001894) has a single LAGLIDADG_ 1 domain, and its amino acid sequence shared high sequence similarity with known LHEs with well-validated activity (Fig. 4). For example, it shared 42% amino acid identity and 56% similarity with I-Ceul, which is encoded by the mitochondrial 23S rRNA gene of the eukaryote *Chlamydomonas eugametos*.

**Fig 4.**
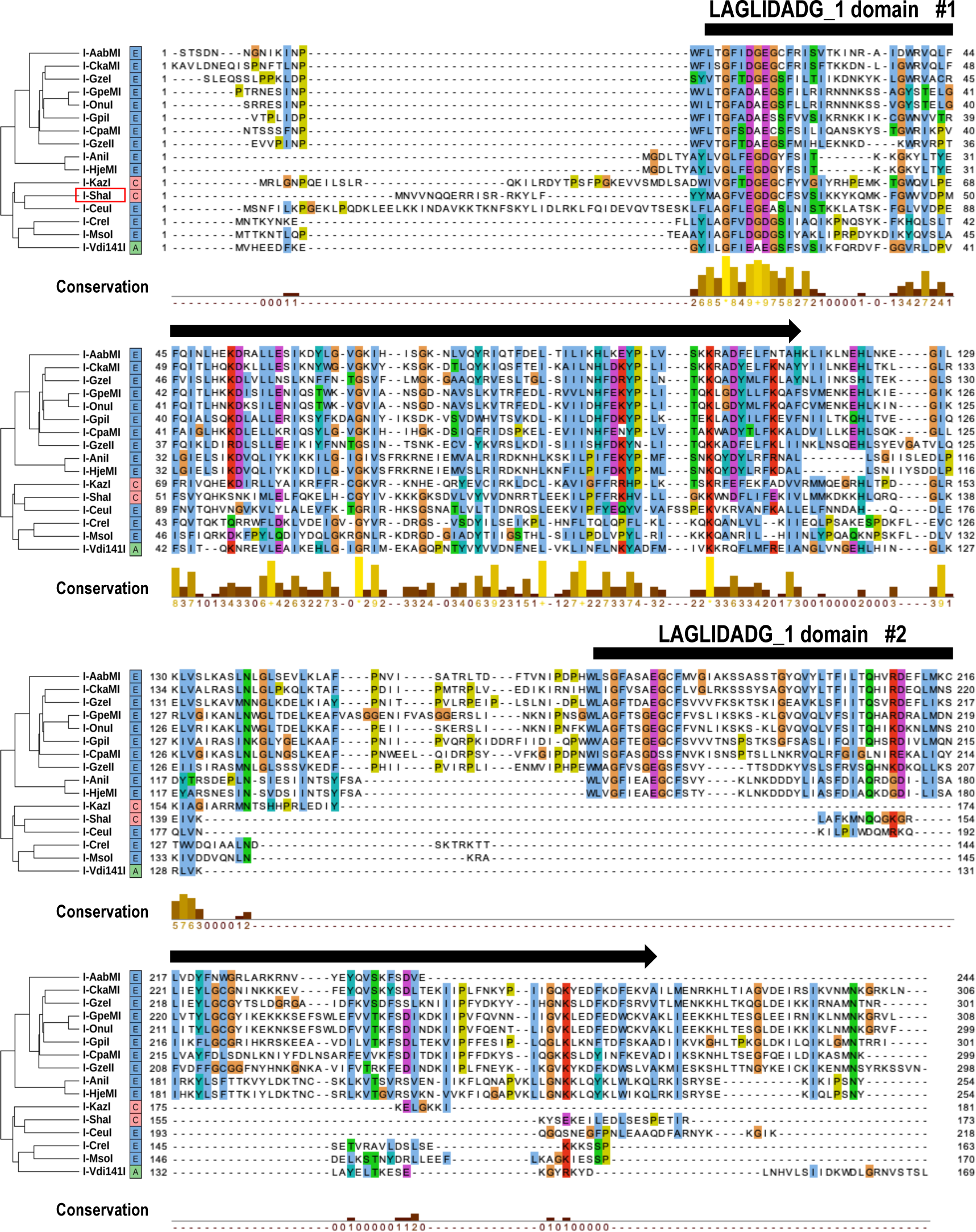
Amino acid sequence alignment of putative LAGLIDADG homing endonucleases from CPR bacteria and other organisms. Amino acid sequences of putative LAGLIDADG homing endonucleases I-ShaI and I-KazI, encoded by ISs in the 23S rRNA genes of CPR bacteria, were aligned with those of 14 LAGLIDADG homing endonucleases with confirmed activity. Boxes on the right of the protein names indicate the taxonomic groups of the host organisms (E: eukaryotes; A: archaea; C: CPR bacteria). Alignment and the average distance tree constructed using the BLOSUM62 algorithm were visualized using Jalview. Consensus amino acids are colored according to the ClustalX color scheme. Conservation scores for each amino acid position are shown as 12 ranks (from 0 to 11) (62). Identical amino acid residues (score 11) are indicated by “*”, and amino acid residues with conserved chemical properties are indicated by “+”. Black arrows above the alignment indicate the LAGLIDADG_1 domain region.

I-ShaI is encoded within the IS of the 23S rRNA gene (positions 2020–2623, 691 nt length, corresponding to 23IS7 in Fig. S1-2), with the ORF positioned at the center of the IS (position 83–604 of the IS) (Fig. 5A). Furthermore, the sequences surrounding the ORF shared nucleotide sequence similarity with group I introns (cmscan with E-value ≤ 1e−4). To validate its endonuclease activity, we first designed an artificial *I-ShaI* gene with optimized codons for its efficient expression in *E. coli* (Fig. S7) and induced its expression in *E. coli* as a His-tagged recombinant enzyme. Recombinant I-ShaI was purified to near homogeneity on SDS-PAGE, by metal affinity column chromatography against the His tag, and anion exchange column chromatography (Fig 5B). The peak elution of recombinant I-ShaI during anion exchange chromatography occurred in fractions #12–#13, and the cleavage activity against plasmid DNA containing the target sequence (Table S6, Fig. S8A) was also at a maximum in these fractions. In other words, a change from supercoiled (SC) to linear (L) plasmid DNA was observed. The molecular weight of the purified protein, calculated from its mobility on SDS-PAGE (approximately 19 kDa), was almost the same as that estimated from the amino acid sequence of the recombinant protein (Fig. 5B). These results indicate that the purified recombinant I-ShaI had activity to cleave the target DNA. The amount of substrate cleaved increased as the amount of added I-ShaI increased, whereas no substrate was cleaved when bovine serum albumin (BSA) was added as the control (Fig. 5C). I-ShaI showed stronger activity at 50–60 °C than at 37 °C, consistent with the fact that the protein was derived from bacteria living in a high-temperature environment (48.5 °C) (Fig. 5D). To examine the substrate specificity of I-ShaI, we tested its activity against different substrate DNAs. The substrate DNA sequences compared were obtained from the genome of *Ca*. Kazanbacteria, another strain of CPR. This genome was assembled from a sample derived from the same environment as the host of I-ShaI, had an IS at the same position in the 23S rRNA gene, and like I-ShaI, contained a single LAGLIDADG_1-domain (designated “I-KazI”; Fig. 4). I-ShaI cleavage activity was also observed (Fig. 5E) when the target DNA substrate was based on the 23S rRNA gene of *Ca*. Kazanbacteria (Fig. S8B). In contrast, there was no cleavage of the control plasmid, in which the target DNA sequence was not inserted. Therefore, I-ShaI has some substrate specificity, but not strict specificity. Like standard HEs belonging to this family, it acts on a wide variety of sequences (37) and can probably induce homing to genomes at a certain phylogenetic distance within the CPR.

**Fig 5.**
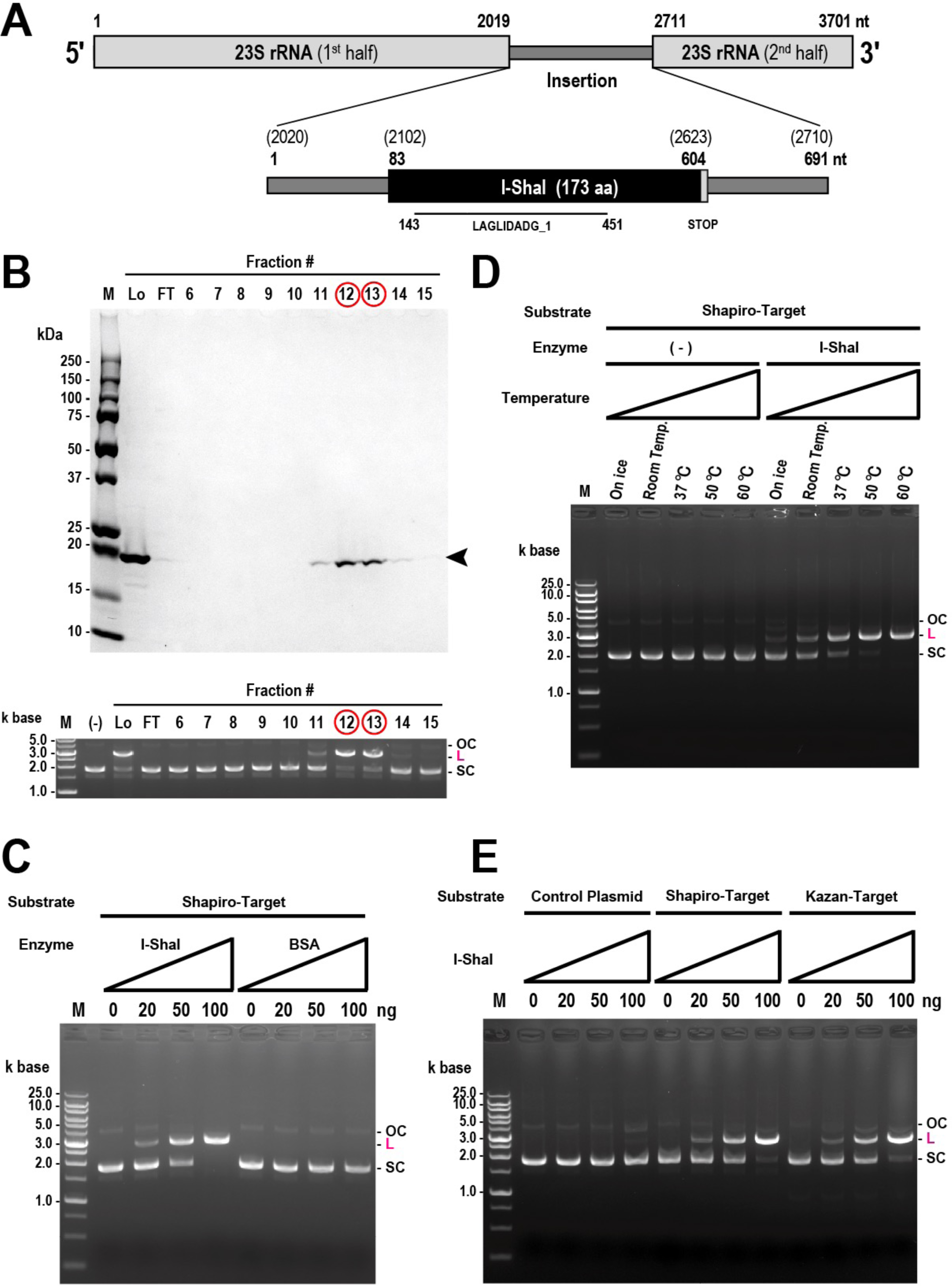
Experimental verification of endonuclease activity of putative LAGLIDADG homing endonuclease I-ShaI. (A) Schematic representation of the putative LAGLIDADG homing endonuclease I-ShaI encoded by the 23S rRNA gene IS in the *Ca*. Shapirobacteria genome (GenBank accession: GCA_003647605.1). Numbers represent nucleotide positions in the gene (upper diagram) or in the IS (lower diagram), and numbers in parentheses in the lower diagram correspond to the positions in the upper diagram. LAGLIDADG_1 domain region in the open reading frame of I-ShaI is indicated by a black bar. (B) Purified I-ShaI has endonuclease activity. (Upper column) SDS-PAGE analysis of I-ShaI purified with HiTrap Heparin HP column chromatography. Aliquots of the fractions from the column were subjected to SDS-PAGE (10%–20% gels), and the gels were stained using Coomassie Brilliant Blue. Lanes M, Lo, and FT indicate molecular weight markers, load fraction, and flow-through fraction, respectively. The arrowhead indicates the positions of purified I-ShaI. Fractions with peak protein elution are indicated by red circles. (Lower column) Endonuclease activity of aliquots of the fractions from the HiTrap Heparin HP column. After the purified enzyme was incubated with the target substrate DNA, the DNA was separated using 0.8% agarose gel electrophoresis and the gel was stained with ethidium bromide. OC, L, and SC indicate open circular, linear, and supercoiled DNA, respectively. Fractions with peak endonuclease activity are indicated by red circles. (C) Dose dependence of I-ShaI endonuclease activity. Bovine serum albumin (BSA) was used as the negative control. (D) Effects of temperature on the endonuclease activity of I-ShaI. (E) Substrate preferences of the endonuclease activity of I-ShaI.

We then analyzed the domain architecture and possible three-dimensional (3D) structure of I-ShaI. Single-domain LHEs are known to function as homodimers (38). To confirm the dimerization of I-ShaI, we used the chemical cross-linker bis(sulfosuccinimidyl) suberate (BS^3^). Upon the addition of BS^3^, a reduction in the band intensity of monomeric I-ShaI (approximately 19 kDa) was observed, accompanied by the appearance of a band at approximately 39 kDa, strongly supporting the formation of homodimeric I-ShaI (Fig. 6A). The 3D structure of I-ShaI, predicted using AlphaFold2, was similar to the structure of a known single-domain LHE (I-CreI; Fig. 6B–D). In known two-domain LAGLIDADG monomers and single-domain LAGLIDADG dimers, the first helix of the core fold acts as the bonding surface between domains (Fig. 6E–F) (37). I-ShaI is also expected to form a dimer in a similar manner, which acts on the target DNA (Fig. 6G).

**Fig 6.**
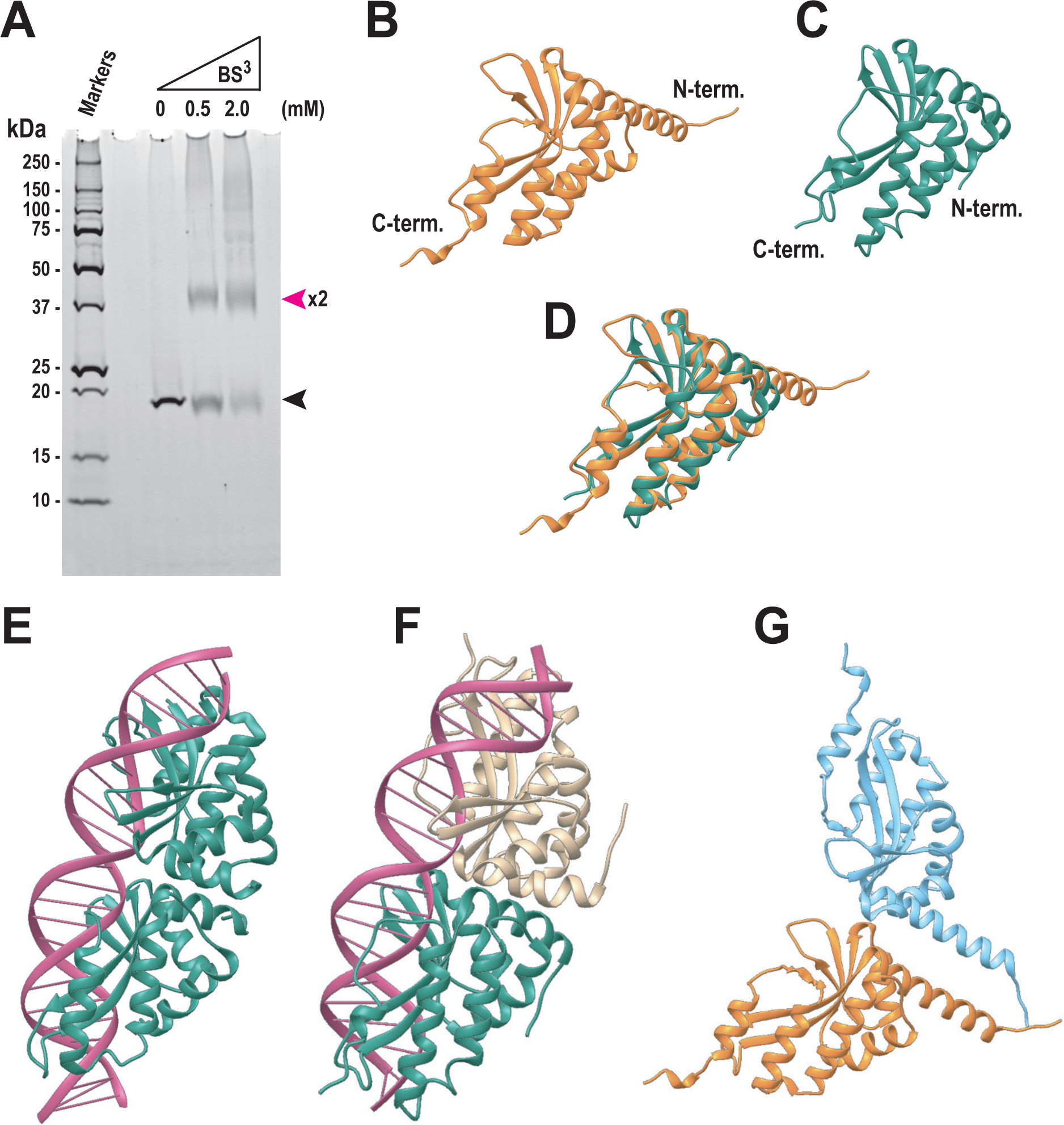
LAGLIDADG homing endonuclease I-ShaI forms a dimer. (A) Chemical cross-linking demonstrated the formation of I-ShaI dimers. A cross-linking assay was performed by treating approximately 500 ng of purified I-ShaI with 0, 0.5, or 2 mM cross-linker bis[sulfosuccinimidyl] suberate (BS^3^). Reactions were stopped and reaction products analyzed using 10%–20% SDS-PAGE. Bands were stained using SYPRO Ruby. Black and pink arrowheads indicate the positions of monomeric and dimeric I-ShaI, respectively. (B) Three-dimensional structure of homing endonuclease I-ShaI monomer, predicted using AlphaFold2. (C) Three-dimensional structure of group I mobile intron endonuclease I-CREI monomer (PDB ID: 1BP7). (D) Superimposition of the structures in panels B and C. (E) Crystal structure of LAGLIDADG meganuclease I-AabMI bound to uncleaved DNA (PDB ID: 4YIT). (F) Three-dimensional structure of homing endonuclease I-CreI dimer complexed with homing-site DNA (PDB ID: 1BP7). (G) Predicted three-dimensional structure of homing endonuclease I-ShaI dimer. The dimeric structure was created by manually placing two predicted monomers.

In summary, we comprehensively investigated the phylogenetic distribution and sequence characteristics of the rRNA ISs in CPR bacteria and analyzed the molecular evolution and biochemical characteristics of the LHEs that are frequently encoded by ISs. Our exhaustive detection and characterization of ISs revealed a much greater abundance of introns and intron-encoded HEs in CPR rRNA genes than has been reported previously in non-CPR bacteria. Self-splicing introns in the 16S rRNA gene, previously reported only in giant sulfur bacteria (32), are abundant in the CPR, and introns in the 23S rRNA gene were observed at diverse sites, exceeding the known intron sites (28–31) in non-CPR bacteria. It remains unclear why such a large number of introns and HEs remain in CPR bacteria. CPR bacteria are thought to have a lifestyle dependent on other organisms based on their reduced genomes, and some lineages have been reported to be the endosymbionts of protists, the episymbionts of other bacteria and archaea, and even predators of the cell contents of other bacteria (16, 22, 24–26, 50). These lifestyles closely associated with other organisms may be responsible for the abundance of ISs, but the absence of ISs in Saccharibacteria, a representative of the CPR symbionts, must be considered. Based on the phylogenetic and positional distributions of ISs encoding LHEs and our phylogenetic analysis of LHEs, the abundance of rRNA ISs in the CPR seems to have resulted from a combination of vertical inheritance, during the branching of the CPR, and horizontal transfer, mainly by homing. The patched distribution patterns of LHE-encoding and LHE-less introns observed at several insertion sites are consistent with the cyclic model of introns proposed previously (44), which involves the gain, saturation, degeneration, loss, and regain of introns by horizontal transfer. Further supporting this model, some of the LHEs encoded at these sites are fragmented into two ORFs or are significantly shortened by the occurrence of a stop codon. However, these may be sequencing or assembly artifacts in the examined CPR genomes, which were derived mainly from metagenomes. Whether the LHEs of CPR bacteria are in the process of spreading by homing or are remnants has been a subject of great interest, and our experimental validation has shown that at least one CPR-derived LHE has target-specific endonuclease activity and dimer-forming ability, indicating that it can potentially drive homing. This is the first study to biochemically characterize the LHEs of CPR bacteria. Our findings provide valuable insights into the evolution and biochemistry of rRNA ISs in CPR bacteria and are an important step forward in our understanding of the CPR.

## MATERIALS AND METHODS

### Dataset

We collected 897 CPR bacterial genomes (69 complete and 828 draft genomes) from the National Center for Biotechnology Information (NCBI) GenBank (https://ftp.ncbi.nlm.nih.gov/genomes/genbank/) (51) in a previous study (21) and extracted their rRNA gene sequences using the method described below. Phylum-level annotations for each genome were assigned based on information from the NCBI Taxonomy Database (52) and previously reported phylogenetic trees (3). We used 1661 complete genomes of non-CPR bacteria annotated as “Reference” or “Representative” in the NCBI Reference Sequence Database (RefSeq; https://ftp.ncbi.nlm.nih.gov/genomes/refseq/; accessed on November 1, 2018) (53) in the analysis of rRNA ISs. To collect the LAGLIDADG proteins from the three domains of life, protein sequences were downloaded from UniProtKB release 2022_04 (October 2022) as a database. The phylogenetic classifications of these proteins were assigned using the NCBI Taxonomy Database.

### Identification of rRNA gene ISs and IS-encoded proteins in CPR bacteria

The rRNA gene sequences were extracted from the CPR bacterial genomes following our previous study (21). In brief, the cmsearch program in the Infernal package (version 1.1.3) (54) was used to search for 16S, 23S, and 5S rRNA genes in the genome sequences (*E*-value threshold: 1e−4). We used the secondary structure models of 5S rRNA (RF00001), 16S rRNA (RF00177), and 23S rRNA (RF02541) registered in the Rfam database (55). Because CPR bacteria often have long ISs within their 16S and 23S rRNA genes, several partial hits were identified. If the partial hits were adjacent on the same scaffold (i.e., the gap between the hits was ≤5000 bases and no hits for other rRNAs were identified in the gap), those hits were considered to be single genes. Partial rRNA gene sequences truncated at the end of the scaffold were excluded from the subsequent analysis. Note that not all of the CPR genomes used here had full-length rRNA genes because many of them were in draft form (see Table S1 for the number of genomes from which any rRNA gene could be extracted). The rRNA genes of non-CPR bacteria were identified according to RefSeq annotations. All copies of rRNA gene sequences were extracted.

The ISs in the rRNA genes were determined based on their alignment with the well-studied 16S, 23S, and 5S rRNA genes of *E. coli* K-12 (NCBI Gene IDs: 948511, 947585, and 947769, respectively). Only ISs of ≥100 bp in length were included in the results because our preliminary search showed that known introns and protein domains were only found in ISs of these lengths. To examine the similarities between the ISs and known group I and II introns, full-length rRNA gene sequences were searched against covariance models in the Rfam database (version 14.6) using the cmsearch program (*E*-value threshold: 1e−5). A hit was considered positive if 25% or more of the hit region in the group I or II intron model overlapped the IS (2).

To detect the ORFs present in the rRNA ISs, we used the getORF program of EMBOSS 6.6.0 (56) to translate the full-length rRNA gene in six frames. In the translation process, genetic code 11 (the standard code for bacteria) was usually applied, with the exception that genetic code 25 was used for Absconditabacteria and Gracilibacteria (57). IS-encoded ORFs were defined as ORFs with a length of ≥10 amino acids that overlapped the rRNA IS by ≥80%. These IS-encoded ORFs were characterized using the domain search described below (see “Protein domain search”), and sequences containing any of the domains LAGLIDADG_1, LAGLIDADG_2, or LAGLIDADG_3 were considered LAGLIDADG proteins.

### Identification of LAGLIDADG proteins

To comprehensively identify the LAGLIDADG proteins in the UniProtKB database, a sequence similarity search was performed using BLASTP (version 2.9.0+) with an E-value threshold of 1e−5. Twenty-three LHEs (see Table S5) obtained from the LAGLIDADG Homing Endonuclease Database and Engineering Server (LAHEDES) (58) were used.

### Protein domain search

Protein domain searches were performed using the hmmscan program in the HMMER 3.3.1 package (59). Using the amino acid sequence of each protein as the query, the HMM profiles of Pfam-A version 34.0 (46), a database of known protein domain families, were searched using an E-value threshold of 1e−4.

### Multiple-sequence alignment analysis

The amino acid sequences of multiple LAGLIDADG proteins were aligned with MAFFT L-INS-i (v7.475) (60). The alignment was visualized using Jalview version 2.10.3 (61). The conservation scores for each amino acid position, calculated according to C. D. Livingstone and G. J. Barton (62), are shown as 12 ranks (from 0 to 11). Identical amino acid residues (score 11) are indicated by “*”, and amino acid residues with conserved chemical properties are indicated by “+”. The alignment files created in this step were also used for phylogenetic tree analysis, which is described in the “Phylogenetic analysis” section below.

### Phylogenetic analysis

For the phylogenetic analysis, we used the alignment file described above and removed any gaps using trimAl (version 1.2, -gt 0.4) (63). Phylogenetic trees were constructed using the maximum likelihood method with RAxML (version 8.2.12) (64). We used the PROTGAMMAAUTO option to select the best model and performed 100 bootstrap replications. The tree was rooted using the midpoint rooting method. The interactive Tree Of Life (iTOL) (65) was used to draw the phylogenetic tree and to map information (such as taxonomy and domain architecture) to the tree. To reduce the number of sequences obtained from UniProtKB when we constructed the phylogenetic tree of the LAGLIDADG proteins derived from organisms in three domains of life (Fig. 2), clustering by CD-HIT version 4.8.1 (66) was performed with a similarity threshold of 40%. Representative sequences of each cluster were combined with all the LAGLIDADG protein sequences found in the rRNA ISs and used to calculate the phylogenetic tree.

### Prediction and visualization of protein 3D structures

The 3D structures of the LAGLIDADG endonuclease I-ShaI were predicted using AlphaFold2 (67) via ColabFold (68) with the default parameters. For structural comparison, the structural data for the known nucleases I-CreI (PDB ID: 1BP7) and I-AabMI (PDB ID: 4YIT) were obtained from the RCSB Protein Data Bank (69). The 3D structures were visualized using UCSF Chimera (70).

### Synthesis of artificial gene encoding I-ShaI protein and construction of expression vector

A CPR Shapirobacteria *I-ShaI* gene (DDBJ accession: BR001894) was used to express the recombinant protein in *E. coli*. First, using a web tool (https://www.eurofinsgenomics.jp/jp/service/gsy/orderstart.aspx?type=MyCart) provided by Eurofins Genomics Tokyo, we optimized the nucleotide sequence to match the codon usage of *E. coli* and synthesized an artificial *I-ShaI* gene. This synthetic gene was designed to contain *Nde*I and *Xho*I restriction sites at its 5′- and 3′-termini, respectively, and was subcloned into these sites in the pET-23b expression vector (Novagen, Madison, WI, USA). The resulting pET-I–ShaI vector encoded the I-ShaI protein with a six-histidine (His) tag at its C-terminus (Fig. S7).

### Expression and purification of His-tagged recombinant I-ShaI protein

To express the recombinant I-ShaI protein, *E. coli* strain BL21(DE3)pLysS was transformed using the expression vector pET-I–ShaI. The transformant was precultured in Luria–Bertani (LB) medium containing 50 µg/mL ampicillin for 4 h at 37 °C. The preculture was then transferred to 200 mL of LB medium containing the same concentration of ampicillin, incubated at 30 °C for 4 h, and then at 17 °C for 2 h. Isopropyl β-D-1-thiogalactopyranoside (IPTG; 0.4 mM) was added and the cells were incubated at 17 °C for a further 14 h to overexpress the desired recombinant I-ShaI protein. The cells were harvested by centrifugation (9000 × g for 7 min at 4 °C), and the protein was extracted by sonication (3–4 min) in His-tag-binding buffer (20 mM Tris-HCl [pH 8.0], 500 mM NaCl, 5 mM imidazole, and 0.1% [v/v] NP-40). The insoluble protein was removed by centrifugation (18,000 × g for 10 min at 4 °C) and purified using HisTrap HP column chromatography (Cytiva, Marlborough, MA, USA) and eluted with a linear gradient of imidazole (0–1000 mM) in His-tag-binding buffer using a ÄKTA™ FPLC™ Fast Protein Liquid Chromatograph (Cytiva). The eluted protein peak was collected and dialyzed against buffer D (50 mM Tris-HCl [pH 8.0], 1 mM ethylenediaminetetraacetic acid, 0.02% [v/v] Tween 20, 7 mM 2-mercaptoethanol, and 10% [v/v] glycerol). The sample containing the dialyzed recombinant protein was then applied to a 1 mL HiTrap Heparin HP column (Cytiva) equilibrated with buffer D and eluted with a linear gradient of NaCl (0–2000 mM) in buffer D using the ÄKTA FPLC system (see Fig. 5B).

### Endonuclease assay

To detect endonuclease activity, two sets of chemically synthesized DNA containing the target sequence (Table S6, Fig. S8) were subcloned between the *Eco*RI and *Bam*HI sites in the pBluescript SK(+) vector (Addgene, Watertown, MA, USA) and designated “pBlue-SapiroTarget” and “pBlue-KazanTarget”. The original pBluescript SK(+) vector with no insert was used as a control. The basic reaction was performed in 20 μL of reaction buffer (10 mM Tris-HCl [pH 7.5], 1 mM dithiothreitol, 50 mM NaCl, 10 mM MgCl_2_, 400 ng of target plasmid, and purified recombinant I-ShaI [0–100 ng]). After the sample was incubated at 50 °C for 60 min, we added 1 μL of 10% SDS and incubated it further at room temperature for 5 min. The resulting sample was separated using 0.8% agarose gel electrophoresis, and the gel was stained with ethidium bromide. The reaction products were visualized using the Molecular Imager FX Pro (Bio-Rad, Hercules, CA, USA).

### Chemical cross-linking analysis

The purified I-ShaI (approximately 500 ng) was incubated in phosphate-buffered saline (PBS) containing 50 mM NaCl and 0, 0.5, or 2 mM BS^3^ (Thermo Scientific, Rockford, IL, USA) at room temperature for 30 min. The reactions were stopped by the addition of 1 M Tris-HCl (pH 8.0), and the products were analyzed by electrophoresis on 10%–20% polyacrylamide gel containing SDS. The protein bands were stained using <u>SYPRO Rub</u>y (Invitrogen from Thermo Fisher Scientific, Waltham, MA, USA).

## ACKNOWLEDGMENTS

We thank all the members of the RNA Group at the Institute for Advanced Biosciences of Keio University, Japan, for their insightful discussions. This work was supported, in part, by a JSPS KAKENHI Grant-in-Aid for JSPS Fellows (grant number 21J12231, to M.T.), a JSPS KAKENHI Grant-in-Aid for Challenging Exploratory Research (grant number 22K19340, to A.K.), the Keio University Doctorate Student Grant-in-Aid Program of the Ushioda Memorial Fund, and research funds from the Yamagata Prefectural Government and Tsuruoka City, Japan. The funding bodies played no role in the study design, data collection or analysis, the decision to publish, or the preparation of the manuscript.

## CONFLICT OF INTEREST STATEMENT

The authors declare that they have no conflicts of interest.

## SUPPLEMENTARY INFORMATION

Supplementary material is available for this article: Supplementary Tables S1–S6 and Supplementary Figures S1–S8.

